# From Coarse to Rich: Successive Waves of Visual Perception in Prefrontal Cortex

**DOI:** 10.64898/2026.03.27.714202

**Authors:** Joachim Bellet, Markus Siegel, Stanislas Dehaene, Bechir Jarraya, Theofanis I. Panagiotaropoulos, Timo van Kerkoerle

**Affiliations:** Cognitive Neuroimaging Unit, CEA, INSERM, Université Paris-Saclay, NeuroSpin, Gif-sur-Yvette 91191, France; Department of Neural Dynamics and Magnetoencephalography, Hertie Institute for Clinical Brain Research, University of Tübingen, Germany; Centre for Integrative Neuroscience, University of Tübingen, Germany; MEG Center, University of Tübingen, Germany; German Center for Mental Health (DZPG), Tübingen, Germany; Collège de France, Université Paris-Sciences-Lettres (PSL), 11 Place Marcelin Berthelot, Paris 75005, France; University of Versailles Saint-Quentin-en-Yvelines, Versailles 78000, France; Neuromodulation Unit, Foch Hospital, Suresnes 92150, France; Department of Psychology, National and Kapodistrian University of Athens, Athens, Greece; Centre for Basic Research, Biomedical Research Foundation of the Academy of Athens (BRFAA), Athens, Greece; Donders Institute for Brain, Cognition and Behaviour, Radboud University, 6525 AJ Nijmegen, the Netherlands

## Abstract

The ventrolateral prefrontal cortex (vlPFC) is well known for its involvement in high-level functions such as cognitive control and language. However, vlPFC’s role in visual processing is less clear. Here, we investigated how neuronal ensembles in the vlPFC dynamically encode different types of visual information. Using chronic recording of spiking activity, we investigated vlPFC’s representational geometry in a macaque monkey passively viewing a large set of naturalistic images, and compared this to representations in deep neural networks (DNNs). We found that the vlPFC processes visual information in two stages. First, an “early” response from 50 to 90 ms after stimulus onset encodes the low spatial frequency component of an image. It contains sufficient information to form a coarse estimate of the position and category of a salient object. Then, from 100 ms on, the representational geometry changes and contains much richer information. This late period contains non-categorical information typically present in conscious experiences such as the orientation of a face and natural scenes in the background. The late window also enables sub-category identification, which is boosted by the low spatial category prior. These results suggest that the vlPFC has a dual role in natural vision: first forming fast low-spatial-frequency-based priors shaping feed-forward visual processing, and subsequently maintaining a detailed and rich representation of a visual scene.

**Significance:** What information does the prefrontal cortex represent during natural vision, and how does this relate to conscious experience? Theories of consciousness differ sharply on whether prefrontal activity reflects detailed perceptual content or only high-level, task-related information. Using dense neural recordings during passive viewing of thousands of naturalistic images, we show that the ventrolateral prefrontal cortex processes visual information in two distinct stages: an early, coarse estimate of the visual scene is followed by a richer, high-dimensional representation that includes sub-category identity and other perceptual details. These findings reveal that prefrontal circuits encode more of the content of visual experience than previously assumed, even in the absence of a task.

## Introduction

The prefrontal cortex (PFC) has traditionally been characterized as a critical region for higher-order cognitive functions, including working memory, executive control, and decision-making (Miller & Cohen, 2001). Yet little is known about the role, if any, of PFC in sensory processing. This question is hotly debated because it sits at the crossroads of differing theories of consciousness: some see the PFC as merely handling what is relevant for higher-order thoughts (Boly et al., 2017), while others argue it encodes anything that is accessible to our conscious awareness including our rawest sensations (Brown et al., 2019; Mashour et al., 2020). While some researchers have consistently measured PFC activity linked to conscious perception (Dehaene et al., 2001; Del Cul et al., 2009; Dellert et al., 2021, Gaillard et al., 2009; Lau & Passingham, 2006; Levinson et al., 2021; Li et al., 2014; Lumer & Rees, 1999; Rounis et al., 2010), others have argued that PFC activity might reflect solely the act of reporting an experience rather than the experience itself (Block, 2011; Brascamp et al., 2015; Frassle et al., 2014; Tsuchiya et al., 2015). Even the correlation of PFC’s activity to the conscious content in the absence of an overt report (Kapoor et al., 2020; Panagiotaropoulos et al., 2012) has been suggested to reflect thoughts occurring as a consequence of consciousness-related processes (Block, 2020; but see Panagiotaropoulos et al., 2020; Dellert et al., 2021, Bellet et al., 2022). Critically, the question remains whether the PFC reflects the richness of conscious experience. For instance, a single glance at someone’s face is associated with multiple perceptual dimensions (e.g. gaze direction, expression, skin color, etc.). A recent study reported that although PFC population signals (eCOG, fMRI, MEG) encode the presence of a face, they do not encode face orientation (Ferrante et al., 2025). However, such null findings are difficult to interpret given the apparent salt-and-pepper functional organization of neurons in the PFC which necessitates neural recordings at a high spatial resolution to find potential representations of detailed visual information (Panagiotaropoulos 2024). An earlier study did describe vlPFC neurons tuned to faces viewing in a specific direction (Romanski & Diehl, 2011). If the neurons responding to right-oriented faces and those responding to left-oriented faces are evenly distributed in the vlPFC then their response would average out at the mesoscopic level. Yet interpreting functional selectivity is a difficult task. Head orientation can be confused with low-level retinal information (Ramírez et al., 2014) and the same applies when investigating category selectivity (Rice et al., 2014).

This raises a simple but fundamental question: what exactly is encoded in the PFC during passive vision? The results of two recent studies favor the view that the PFC represents low-level visual features rather than those corresponding to conscious experience. The first study showed that the vlPFC neurons are better driven by large blob-like stimuli than by high-level features (Rose & Ponce, 2024). The second reveals that PFC neurons can discriminate faces from non-face stimuli as early as 40 ms after image onset and that their firing rate is higher for images dominated by low spatial frequencies (LSF; Mergan et al., 2025). Together, these findings provide support for the top-down facilitation theory proposing that the PFC provides early priors based on LSF that shape object recognition in the ventral stream (Bar, 2003). While such rapid LSF processing likely reflects non-conscious mechanisms, these observations do not preclude that the PFC may subsequently encode the detailed information corresponding to conscious experience. Indeed our own work revealed the vlPFC visual codes are dynamic, leaving open the possibility that finer image details are represented after the early LSF information (Bellet et al., 2022). Thus, the vlPFC may have the dual role of shaping visual processing via its top-down connections, and simultaneously represent the many dimensions of conscious experience.

To address these questions, we probed vlPFC representations during passive vision using multi-site recording of spiking activity in the vlPFC of a macaque monkey viewing a large, parametrically controlled image set. We find that 50 to 90 ms after stimulus onset, the vlPFC already contains category information solely carried by the LSF component of the images. After 100 ms, the geometry reorganizes and becomes markedly richer. First, sub-category within the same category can now be separated from each other and this recognition benefits from the LSF category priors. Second, for face stimuli, the population geometry reflects their size, position, and gaze orientation. Finally, we find that even the landscape in the background of the salient object is represented by vlPFC. These results demonstrate that, even without an active task, vlPFC neuronal ensembles can generate rapid LSF-based top-down facilitation of object recognition on a millisecond scale and subsequently represent the many dimensions that compose conscious vision.

## Results

### Dynamics of representational geometry

We started our investigation by measuring at what time visual information reaches the vlPFC and testing whether the functional selectivity of the vlPFC population is constant or if different features are processed at different times from stimulus onset. To this end, we acquired spiking activity with a multielectrode array chronically implanted in the left vlPFC of a macaque monkey who was passively viewing a large set of visual stimuli (Fig. 1*A-B*; see methods for details). The neuronal data aligned to stimulus onset was then split into two independent sets and for each of these folds we measured how dissimilar the neuronal ensemble state was between the different images. The correlation between the representational dissimilarity matrices (RDMs) of the two folds was computed to give a measure of the “geometry consistency” (see methods). Importantly, to ensure that the observed representational geometry was not driven by a subset of images and generalize to novel images, we computed these consistency metrics by resampling the stimulus set into multiple independent batches of images. This measure, computed for each time-bin, revealed that the vlPFC starts to have consistent responses already between 50 and 60 ms after stimulus onset and represents image-specific information until the time of the next stimulus onset at 200 ms (Fig. 1*C*).

**Fig. 1.**
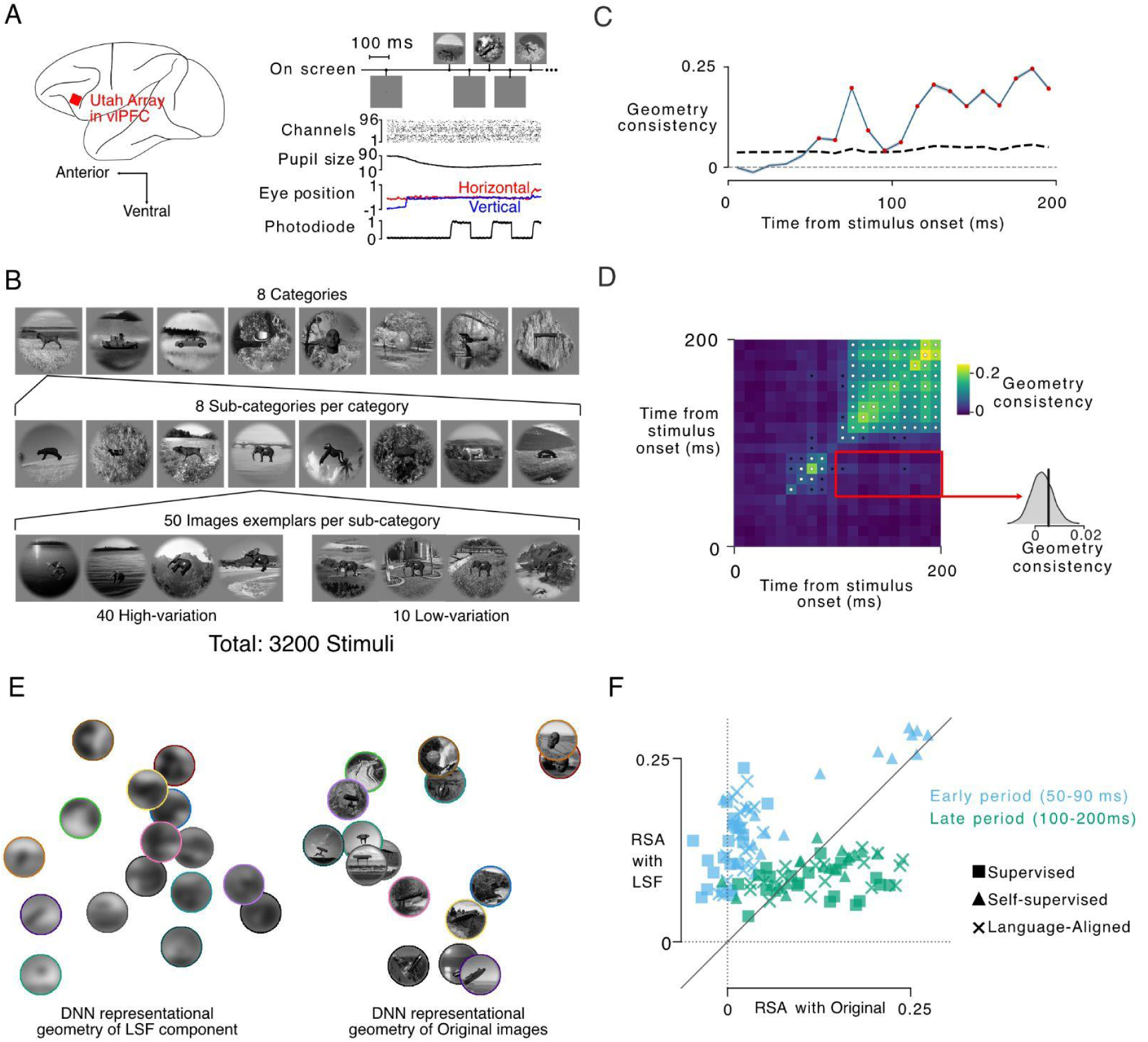
The vlPFC processes two different spatial frequency contents sequentially. (A) Experimental design. Left: Schematic location of the Utah array implanted in the vlPFC. Right: Timeline of an example trial together with recorded signals. From top to bottom: Stimuli presented successively on screen. Raster of spikes from all 96 channels, pupil size (plotted as percentile), horizontal and vertical eye position (degrees of visual angle), and the photodiode signal used to timestamp. (B) Stimulus set structure. Images were drawn from 8 high-level categories (top row). Each category contained 8 distinct sub-categories (middle row). For every sub-category, 50 images were rendered (bottom): 40 “high-variation” images in which the object position, size, and 3D rotation varied together with the natural background, and 10 “low-variation” images in which only the background changed while the object remained centered and identically posed. In total: 3200 stimuli (2560 high-variation; 640 low-variation). (C) Time-resolved geometry consistency between odd- and even-trial neural RDMs. Blue line shows observed Spearman correlations; red dots indicate FDR-significant time-bins; black dashed line, mean surrogate value; grey dashed line, 95% surrogate envelope. (D) Cross-temporal geometry-consistency matrix showing correlations (Spearman r) between neural RDMs at all pairs of time points. White dots show FDR-significant pairs of time-bins. Black dots show pairs of time-bins with p<0.05 but not surviving the control for multiple comparisons. Inset: the average of cross-time correlation between all the time-bins of the early and the late period (vertical line) is not significantly higher than chance (null distribution). (E) Two-dimensional metric-MDS of DNN representational geometry of an example set of images. Left: when using low-spatial-frequency (LSF) images as input to the DNNs. Right: when using the same stimuli but in their Original version. (F) Scatterplot of correlation between neural RDMs and deep neural network DNN RDMs computed from either LSF images (y-axis) or Original images (x-axis). The diagonal indicates equal correlation with Original-image and LSF-image representations. Note: All face stimuli shown are computer-generated 3D models from the Majaj et al. (2015) dataset and do not depict real human individuals.

Next, we probed the temporal stability of representations by computing the geometry consistency between every pair of time points (Fig. 1*D*). This revealed that the representational geometry starting 50 ms after the stimulus onset lasts until 90 ms and is then replaced by a completely different geometry from 100 ms to 200 ms. In the following we will reference these two periods as the “early” phase (50 to 90 ms) and the “late” phase (100 to 200 ms) respectively. And because they are dissociable, all following analyses treat these two windows separately.

We wondered whether the biphasic response geometries stemmed from the early period processing exclusively the low-spatial-frequency (LSF) content while the late period accesses finer visual details. To test whether visual information in the early and late phases differs in its spatial frequency content, we computed representational similarity analysis (RSA) between neural activity and 60 different deep neural networks (DNN) exposed either to the same images as the monkey did or to blurred versions of these images. We included this extensive panel (encompassing 20 supervised models, 20 self-supervised models, and 20 language-aligned vision encoders) to ensure our findings generalize across varied architectures and to test whether specific training regimes align better with vlPFC representations. A two dimensional embedding of the representational distances in the penultimate layer of DNNs revealed how the representational geometry can drastically differ when only the LSF content or the full image (Fig. 1*E*) was used as an input to the network. While vlPFC representations in both time windows aligned significantly above chance with DNN features across all image conditions (all permutation test q < 0.001), we observed a biphasic shift in the preferred spatial frequency content. The representational geometries in the early phase of vlPFC responses correlated significantly more with the representations of DNNs processing the blurred version of the images than the original images (Fig. 1*F*; early mean RSA: Original = 0.043 ± 0.010, low-pass = 0.155 ± 0.008; Wilcoxon test: p = 1.71 x 10^-11^). We noted that self-supervised networks, that are often trained at matching blurred images with unaltered counterparts, tended to align more strongly with early vlPFC activity compared to both supervised (FDR-corrected q = 3.7 x 10^-3^) and language-aligned models (q = 2.1 x 10^-2^). In contrast, the late phase showed the opposite pattern: neural representations correlated more strongly with Original images (Fig. 1*F*; late mean RSA: Original = 0.121 ± 0.008, low-pass = 0.090 ± 0.003; Wilcoxon test: p = 5.26 x 10^-4^) with no significant differences found between training regimes (Kruskal-Wallis test: H(2) = 0.439, p = 0.803).

These results support the hypothesis that the early neural representation in vlPFC primarily encodes low spatial frequency content, whereas the later representation integrates higher spatial frequency details. This raised a critical question: does the early phase still contain meaningful information that could contribute to object categorization?

### Low spatial frequency content contains a category prior also present in vlPFC representations

Is there a functional value for the LSF component to reach vlPFC in the early phase (50 to 90 ms) or is this response epiphenomenal? Since the early vlPFC responses were similar to that of the DNNs processing coarse images, it is likely that DNNs have learned to extract information from the low frequency spatial component of images during their training objectives. In line with the theory of top-down facilitation of object recognition (Bar, 2003), we tested whether the category of an object can be inferred from the LSF component only. We ran logistic regressions to decode the category using the penultimate-layer features from the DNNs fed with the low-pass versions of the stimuli. Importantly we designed the cross-validation strategies of the classifiers such that only sub-category-agnostic category information could be extracted (Fig. 2A). This ensures that the classifiers could not use sub-category-specific features as shortcuts and that the measured information generalizes to novel stimuli. We found that the penultimate layers of the DNNs processing the LSF images contained significant information discriminating between broad categories (Fig. 2 B). This information was not dependent on specific training regimes (Fig S2).

**Fig. 2.**
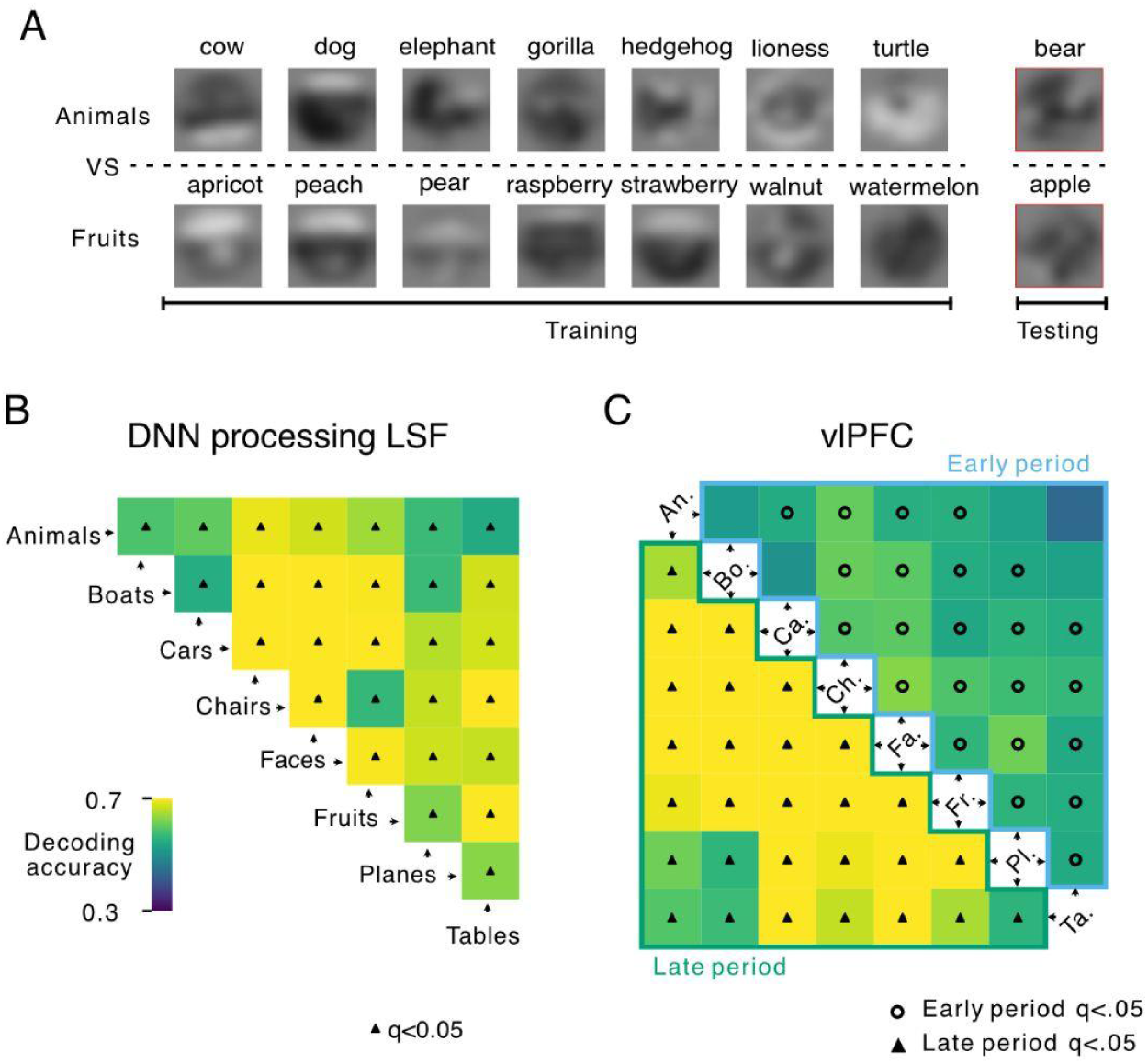
Category information is present in the low spatial frequency component of images and in both phases of the vlPFC response. (A) Schematic of the leave-one-sub-category-out cross-validation strategy for decoding category membership. To ensure that classification is not driven by sub-category-specific features, the classifier is trained to discriminate between two categories (e.g., Animals vs. Fruits) using all but one sub-category per category (e.g., excluding ’bear’ and ’apple’). The accuracy is then evaluated on the held-out pair of sub-categories. (B) Pairwise category-decoding accuracy derived from DNN features of the LSF images. Each cell shows linear-classifier accuracy for distinguishing two categories; black triangles mark decoding significantly above chance (q<0.05, FDR-corrected). (C) Pairwise category-decoding accuracy based for the early (50 to 90 ms, light blue outline) and late (100 to 200 ms, dark green outline) response windows vlPFC of activity. Circles indicate significantly above-chance category discriminability in the early window (q<0.05); triangles indicate significance in the late window.

We next asked if the category information is indeed present in the vlPFC’s response. We employed the same decoding strategies as with the DNN embedding but using the spiking responses of the vlPFC separately in the early (50 to 90 ms) and the late (100 to 200 ms) periods. Most pairs of categories (24/28) and all pairs (28/28) could be discriminated against during the early and late vlPFC responses respectively (Fig. 2*C*). This confirmed that the fast responses of the vlPFC carries a signal that can be used as a prior for fast top-down modulations.

### Low spatial frequency category prior boosts sub-category information in vlPFC

We next tested whether the neural activity in vlPFC contains enough detailed information to distinguish between the different sub-categories within a category, e.g. between a lion and an elephant. For each sub-category, we defined a prototype as the mean neural response across its low-variation exemplars (Fig. 1B). We then defined distance to prototype (DP) as the mean squared euclidean distance between responses to high-variation exemplars and that prototype (see Methods). We tested if the average DP measure was smaller when using images from the same sub-category as the prototype compared to images from a different sub-category yet from the same category (Fig. 3*A*). We found that during the late period, the average DP is smaller than chance (p = 0.0124, q = 0.0248; Fig. 3*B*) which is a signature of object recognition. In contrast, the average DP in the early period did not differ from a chance distribution generated by randomly shuffling exemplars within the same category (p = 0.1477, q = 0.1477). This indicates that while the broad category can be guessed in the early period, only the late period carries information about the sub-category.

**Fig. 3.**
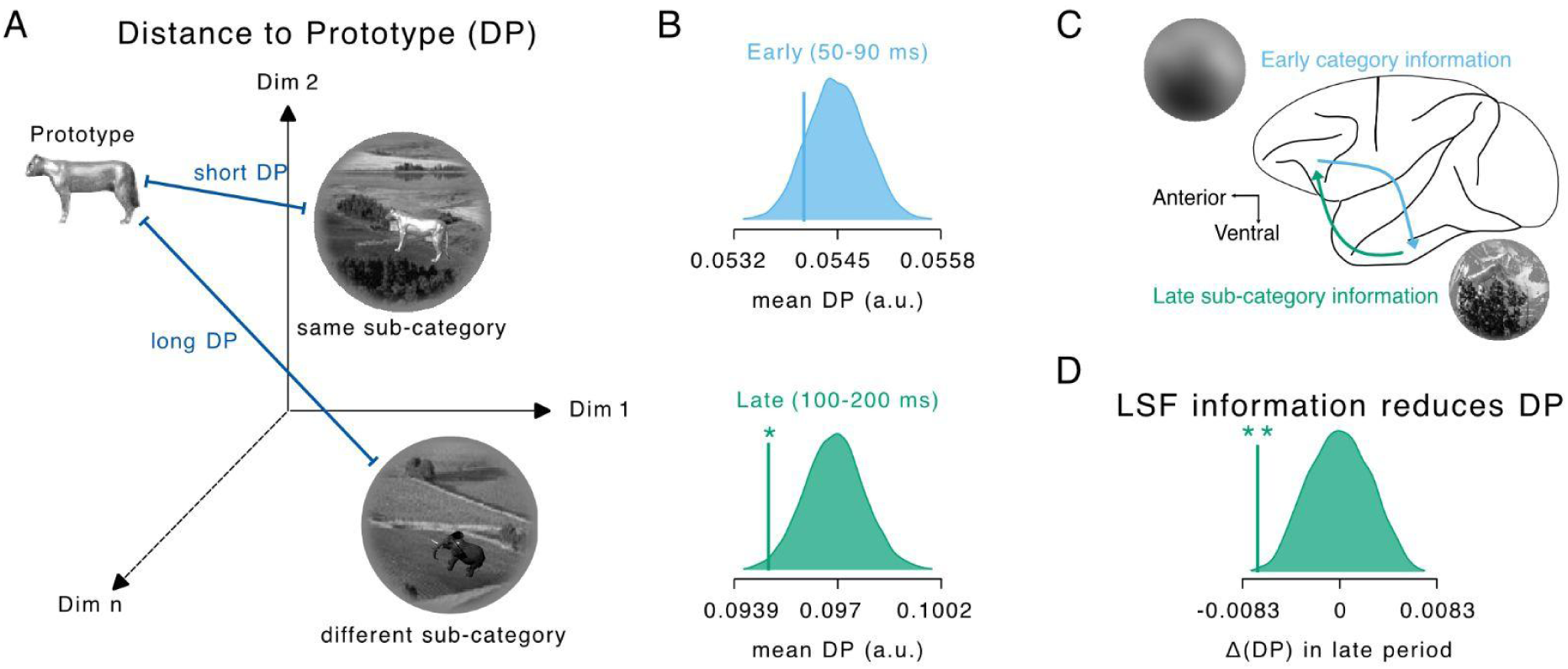
LSF-based priors enhance late-phase sub-category information in vlPFC. (A) Distance-to-prototype (DP) framework. Schematic illustrating the quantification of sub-category information in neuronal state space. Proximity to a sub-category prototype serves as a metric for object recognition. (B) Early vs. late sub-category discriminability. Vertical lines are mean DP values for same-sub-category images during the early (50 to 90 ms) and late (100 to 200 ms) response windows. The null distribution is obtained by surrogate averages using responses to images from other sub-categories but from the same broad category. Only the late responses depart from the null distribution, indicating sub-category specificity. (C) Category-prior gain hypothesis. Schematic of how coarse LSF inputs may provide an early estimate of object category, which could sharpen late-phase sub-category representations. In this illustration the more the LSF contains information about the broad category (here category “animal”), the closer the late vlPFC representation will be to the prototype representation (here “lioness” prototype) (D) Empirical category-prior effect. Difference in DP between images with high vs. low LSF-based category probability (as estimated by DNNs). Negative values indicate that LSF-derived category priors enhance late-phase discriminability. The vertical line marks the experimentally observed difference in mean DP, which is compared against a null distribution (shaded area) generated by shuffling the high/low LSF-information group attributions.

According to the top-down facilitation hypothesis by Bar, it would be expected that information about the broad category in the early period provides a prior that can help define the more detailed sub-category in the late period (Bar, 2003). For instance, if the early wave of activity contains evidence that there is an animal, then its sub-category (e.g. a dog) can be better identified during the late wave of activity. To test this, we asked whether the broad category information carried by LSF content could improve sub-category identification during the late vlPFC response (Fig. 3*C*). We quantified these priors using the continuous outputs of classifiers trained on DNN features processing LSF images and computed, within each sub-category, the difference in DP between the high vs. low prior subsets (median split). DP was significantly smaller when LSF evidence favored the correct category (p = 0.0017, q = 0.0034; Fig. 3*D*). This facilitation was not a consequence of the objects’ saliency with respect to the landscape in the background because the object position information accessible from the blurred image (Fig. S1*A* and S1*B*) and also present in the vlPFC representation (Fig. S1*C* and Fig. 4*A*) does not influence object identification (p = 0.2685, q = 0.2685; Fig. S1*D*).

**Fig. 4:**
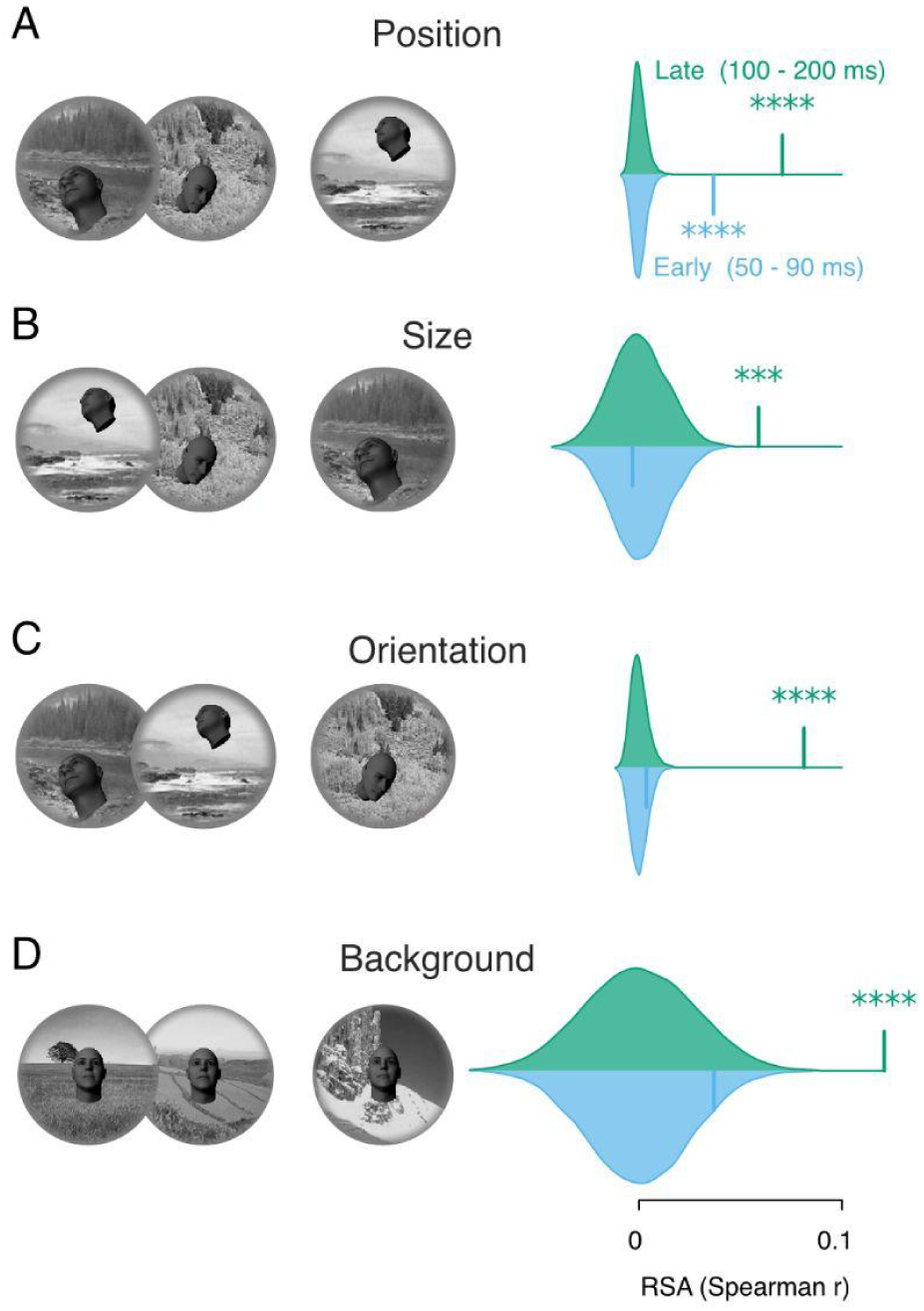
The major dimensions that characterize the consciously visible scene are present in vlPFC representational geometry. (A) Representational similarity of vlPFC geometry with head position. Left: illustration of the tested geometry. Right, observed RSA in the early and late windows of vlPFC responses. Vertical lines indicate measured RSA values and gaussian distributions represent associated chance levels. (B) Same as A. but measuring if the vlPFC geometry reflects head size. (C) Same as A. but for head orientation. (D). Same as A. but measuring if vlPFC senses the natural background behind the foreground object (geometry extracted from language-aligned image models). In all panels stars indicate FDR-corrected significance (**** q < 10⁻⁴, *** q < 10⁻³, ** q < 10⁻², * q < 0.05). Note: All face stimuli shown are computer-generated 3D models from the Majaj et al. (2015) dataset and do not depict real human individuals.

Our result suggests that even in passive viewing conditions, the vlPFC forms sub-category representations that benefit from the early LSF-based category prior.

### Conscious dimensions are present in the vlPFC representational geometry

If the late vlPFC supports fine object identification can it also represent perceptual dimensions that are orthogonal to the object category? This is an important prediction of theories of consciousness proposing a pivotal role of the PFC in constructing subjective experience: all the perceptual dimensions that are consciously visible in a visual scene should be reflected in the PFC activity. To explore this, we examined how vlPFC representations capture perceptually salient features that our subject may also perceive: the position, size and orientation of face stimuli and the naturalist background behind the 3D object. We found that the early period geometry correlated only with the position geometry (Fig. 4*A*), consistent with the cross-category position decoding that we reported above (Fig. S1). In contrast, the late period geometry correlated significantly with the position (Fig. 4*A*), the size (Fig. 4*B*) and the orientation of faces (Fig. 4*C*). In addition, we found that the late geometry reflects also the information about the background landscape as measured by embedding of language-aligned DNNs processing in-painted versions of the images where the central object was artificially removed (Fig. 4*D*).

These results demonstrate that the late vlPFC response contains many category-orthogonal representations typically constitutive of the phenomenal experience of consciousness.

## Discussion

In the present study, we showed that the vlPFC population responds to natural images in two temporally distinct phases. Within 50–90 ms after stimulus onset, the representational geometry aligns with the low-spatial-frequency (LSF) structure of the images and carries coarse information about object position and broad category. From 100 ms onwards, the geometry reorganizes and becomes richer: objects within the same category can be distinguished from each other, and the vlPFC representational space reflects face position, size and orientation, as well as information about the background scene.

Growing evidence shows that the brain takes advantage of early LSF information to guide later finer-grain visual processing. In mice, primary visual cortex neurons rapidly shift their preferred spatial frequency from low to high, reducing redundancy and leading to a more efficient neural representation of natural scenes (Skyberg et al., 2022). In macaque inferotemporal cortex, neurons are first tuned for face detection before rapidly switching around 100 ms to finer face-specific features (Shi et al., 2023). Moshe Bar made the bolder claim that the PFC shortcuts the well established cortical hierarchy and uses an early extraction of LSF information to facilitate visual processing in a top-down fashion (Bar, 2003). Until now evidence in favor of this hypothesis was based on source-reconstructed MEG signals and slow fMRI responses during active recognition tasks (Bar et al., 2006; Musel et al., 2014; Petras et al., 2019). Recent direct recordings of spiking activity in macaque PFC gave more credit to this hypothesis by showing that stimulus identity can be decoded as early as 60 ms (Bellet et al., 2022) and that PFC can respond to faces stimuli faster than IT with more vigorous responses for low spatial frequency stimuli (Mergan et al., 2025).

The present results refine this picture in two ways. First, we show that only the early response in the period 50 to 90 ms reflects LSF information and that it suffices to distinguish the coarse category of picture which is being presented (e.g. animals; it is important that this information is not only restricted to faces). Second, we show that the LSF category prior can boost sub-category information in the later period just like the top-down facilitation theory predicts. Importantly, as the LSF category information was extracted from DNN features, this prediction signal cannot be attributed to trial-to-trial fluctuations in signal to noise or to idiosyncrasies in the response strength of the recorded neuronal population in response to different stimuli. Moreover, since each object was left out in the training phase of the classification procedure, features associated with enhanced object identification are those being common to all objects in the same category. This supports the idea of a push-pull mechanism where a first coarse hypothesis about “what kind of thing is there” helps select the finer features needed to distinguish specific objects. The fact that this effect is restricted to non-spatial features is in line with the crucial role of the vlPFC in feature-based attention and not spatial attention (Bichot et al. 2019).

The fact that category information reaches the penultimate layers of the DNN processing with blurred images, indicates that vision models also learn to use LSF information to optimize their objectives. It has indeed been demonstrated that foundational vision models are highly sensitive to LSF and that this property enhances their robustness to adversarial attacks (Li et al., 2023). However, unlike the brain, these models achieve their performance through deep feedforward architectures trained on vast datasets, whereas biological systems reach comparable capacities from noisy retinal inputs and far fewer synaptic steps, perhaps compensating shallow network depth with recurrent processing (Kar et al., 2019; Kietzmann et al., 2019). While feedforward architectures are still gold standard in machine vision, recent work reveals that recurrent models outperform feedforward counterparts (Greco et al., 2025).

The late geometry captures face position, size and orientation, and correlates with background scene structure when the foreground object is removed and the background is reconstructed. These dimensions are canonical constituents of visual experience and extend beyond what is strictly needed to categorize objects. Together, these results suggest that during passive viewing the vlPFC maintains a scene-level representation that could support conscious access. Importantly, our results are at odds with the conclusion of the Cogitate consortium (Ferrante et al., 2025) who interpreted the absence of significant face orientation decoding in PFC as evidence against a key prediction of the GNWT. Given the salt-and-pepper organization of neuron selectivity in the PFC, such codes are unlikely to be detectable with the coarse spatial sampling of fMRI, ECoG and MEG used by the Cogitate consortium. This is probably for the same reason that, despite PFC’s well established role in working memory, fMRI-based decoding studies often reveal working memory correlates in posterior parts of the brain but not in the PFC (Lara & Wallis, 2015).

We note several limitations of the present study. First, this is a single-animal study which merely provides an existence proof that the vlPFC can exhibit all the properties reported above. Such a study is informative and fully legitimate given the emphasis on the “3R rule” in non-human primate research (replace, reduce, refine). However, the question does remain whether the findings are universal and how they vary across individuals. Second, our analyses are based on a limited subset of the vlPFC population and each stimulus response is averaged across multiple presentations. Whether all aspects of conscious experience are represented simultaneously in a single trial, or whether the PFC selects between these perceptual dimensions on a trial by trial basis, remains an open issue which could be overcome in future experiments by using high density recording, such as large scale calcium imaging or neuropixel probes, to measure enough information at the level of a single trial. Finally, although our results are consistent with PFC contributions to conscious representation, the use of a passive no-report design, while advantageous to avoid additional decision-making processes, cannot test whether the measured dimensions were actually consciously perceived by the subject.

Our study leaves the question of how the vlPFC receives such rapid visual information. The classic ventral stream hierarchy involves many synaptic relays from the retina, but one possibility is that fast information is being conveyed by the very first spike at each synaptic relay rather than by a rate code (Thorpe et al., 1996, 2001). The early phase of processing might predominantly be driven by the dorsal stream, where latencies are shorter, e.g. because of skipping the superficial layers of V1 (Schmolesky et al., 1998; Lamme & Roelfsema 2000). Alternatively (or additionally), the fast LSF information may use subcortical routes from the retina to the vlPFC. Indeed, the retina projects to the superior colliculus which in turns projects to high level thalamic structures directly connected with the vlPFC such as the medio-dorsal thalamus (Sommer and Wurtz, 2006) and the pulvinar (Romanski et al., 1997).

In conclusion, this study provides evidence for a dual role of vlPFC during vision composed of two phases. The first phase accesses coarse image information to form quick priors which are facilitating object information. The object information itself is encoded in a later phase and is accompanied by the representation of perceptual attributes which are typically constituting the phenomenal experience of vision. Our findings confirm that vlPFC encodes the details of a visual scene, which in turn supports the possibility of a crucial role for vlPFC in the emergence of phenomenal consciousness.

## Experimental Methods

### Animal model and ethical considerations

Our study necessitates the direct measurement of spiking activity to investigate the computations of ensembles of neurons in the vlPFC. This area does not have a clear homologue in non-primate animals (Preuss & Wise, 2022), justifying the choice of the macaque as an animal model. The ethical burden of such a study forces us to conduct the experiment in as few animals as possible. As recently demonstrated by Fries and Maris (Fries & Maris, 2022), using one animal is sufficient for an “existence proof”, meaning to test whether a property can exist. The probability of observing the same effects in all animals is informative when one can show that the effect can be inferred to occur in at least 50% of the population and this would require measuring the same effect in at least 5 animals. We therefore chose to perform the experiment in one animal only, leaving open the question as to whether the effects reported here are subject-specific or universal to all primates.

### Subject

One male macaque monkey (Macaca mulatta, 19 years old and 9 kg) participated in this study. He was pair-housed and his water access was controlled during the experimental period which lasted 24 days. The monkey was re-employed after having participated in previous experiments in the laboratory. As he was already implanted with a titanium headpost and a Utah array placed in the vlPFC, no additional surgery was required. Animal handling, housing, and experimental procedures adhered to the guidelines set forth by the European Convention (86/609/EEC) and the National Institutes of Health’s Guide for the Care and Use of Laboratory Animals. The study received approval from the CEA Ethical Committee (CETEA protocol number A18_028).

### Task Design

The monkey was trained to sit in a primate chair and to fixate stimuli displayed on a monitor. Crucially, the monkey never learned active tasks requiring to report what is displayed on the screen and was naive to the image set. Therefore, the results presented in this paper are not confounded with task learning or intention to report any image properties. The monkey’s task was to fixate his gaze at the center of the screen for a few seconds while a series of images were presented in rapid succession. At the end of such a trial, the monkey received a drop of a liquid reward consisting of a mixture of apple juice and water.

No reward was given if the monkey blinked or looked more than 5 degrees of visual angles (DVA) away from the center of the screen before the end of a trial. The number of images presented in a trial could be changed during a recording session to be adapted to the monkey’s motivation. The ratio of apple juice over water was also manipulated by the experimenter and increased toward the end of a recording session, causing a renewed interest of the monkey for executing additional trials. Between any two trials, a fixation spot subtending 0.1 degrees of visual angle was displayed at the center of the screen. The monkey could choose when to start a trial by initiating fixation and maintaining it for 300ms. After this delay, a random subset (median: 7, min: 1, max: 11) of the 3200 images was presented. Each image subtended 8 DVA. The stimulus onset asynchrony was 200 ms with each image being presented for 100 ms and followed by a 100 ms gray screen. The liquid reward was delivered at a random time between 100 ms and 300 ms after the offset of the last image.

### Stimulus set

The image dataset was developed by Majaj and colleagues to study object recognition by minimizing confounds with low-level image features (Majaj et al., 2015). Briefly, each image featured a 3D object placed at a random position and rotation in front of a naturalistic background. Crucially, no two images had the same background and the background was unrelated to the category of the object. Each of the 3200 stimuli belonged to one of the 8 following categories : airplanes, cars, chairs, faces, animals, boats, fruits, and tables. Each of the 8 categories contained 8 representative sub-categories, e.g. bears, lions, elephants etc. for animals. Thus, the dataset contains 64 sub-categories and for each sub-category it contains 50 unique images, which we call ‘exemplars’. Out of the 50 exemplars of a sub-category, 40 images consisted of high variation in the object position, size and rotation while 10 images presented the object at the center, in the exact same position but with varying backgrounds. We call these two different subsets “high-variation” and “low-variation” respectively. See Fig. 1*B* for an overview of the dataset structure.

### Chronic recording of spiking activity

The monkey had been implanted in the vlPFC with a 96 channels UTAH array (Blackrock Neurotech) targeting the inferior convexity (area 45a). We placed the array more towards the lateral side of the vlPFC as this region is known to receive more input from anterior IT, and therefore likely corresponds to later stages of visual processing (Chavis & Pandya 1976; Pandya & Yeterian 1985; Petrides & Pandya 1988; Webster et al. 1994). The wide-band electrophysiological signal was first band-passed between 500 and 6000 Hz using a zero-lag finite impulse response of order 200. The time of spikes was obtained whenever this filtered signal crossed a threshold defined by 4 standard deviations. Because spikes themselves increase the variance of the filtered signal relative to a normal distribution, the robust estimate of the standard deviation was determined using the formula 𝑆𝐷 ≈ 𝑚𝑒𝑑(|𝑋 − 𝑚𝑒𝑑(𝑋)|)/0. 6745 where X is the filtered signal.

### Eye tracking

The eye position of the animals was monitored at 500 Hz, using an infra-red camera (GS3-U3-41C6NIR-C, FLIR Integrated Imaging Solutions, Inc.) and free eye tracking software (iRecHS2, Human Informatics Research Institute, National Institute of Advanced Industrial Science and Technology; Matsuda et al., 2017).

### Spike time alignment

The following steps were applied for each of the 16 recording sessions. For each stimulus presentation, we extracted the precise spike time responses relative to stimulus onset using the photodiode signal. We computed the spike count in bins of 10 ms in the period 0 to 200 ms relative to stimulus onset. A stimulus presentation was excluded from further analysis if 1) the eye position deviated by more than 2 DVA relative to image center, or 2) if the same stimulus had appeared in either of the two preceding trials, or 3) if the mean firing rate during stimulus presentation fell outside the 0.1–99.9 percentile range of its session. Within each session we first averaged repetitions of the same image, yielding one 96 × 20 matrix per stimulus. We then subtracted, for every channel, the session-wide mean firing rate across all stimuli and across all time-bins to remove day-to-day baseline shifts. Finally, we split the dataset by the chronological session index: sessions 0, 2, 4 … contributed to the even half and sessions 1, 3, 5 … to the odd half. For each of the 3200 stimuli, we averaged all valid, baseline-shifted trials across recording sessions within its corresponding half.

### Analyses of representational geometry dynamics

To determine when vlPFC population activity becomes reliable enough to encode visual information we measured how similar the representational geometries were between two independent halves of the dataset (odd sessions vs. even sessions). For each of the 2560 high-variation stimuli and each 10 ms time-bin, we computed average spike counts in the odd and even halves separately. For every time-bin, we then built representational dissimilarity matrices (RDMs) by taking euclidean distances between the population responses to all pairs of stimuli. Geometry consistency was defined as the Spearman correlation between the odd-session and even-session RDMs. Computing a single 2560 x 2560 RDM per time-bin would have required storing 3,275,520 unique pairwise distances per bin (2560 x 2559 / 2) and repeatedly correlating very large matrices, which is computationally demanding and memory intensive. The 2560 stimuli were therefore divided into 160 batches of 16, and a round-robin scheduling procedure generated ten different batch configurations. Each run produced different pairings of stimuli, so that many unique stimulus pairs contributed across the ten runs. For each time-bin and batch, we computed the Spearman correlation between the odd-and even-session RDMs, and then averaged these correlations across batches and across runs. This “average of within-batch correlations” provides a stable summary of geometry consistency that is not driven by any particular grouping of stimuli. We also asked whether the geometry observed at one time point resembled the geometry at another time point. For this purpose, we computed cross-temporal geometry consistency. Instead of correlating odd and even RDMs only within the same time-bin, we computed correlations for every pair of bins across the full 0 to 200 ms window. This produced a time-by-time matrix for each run, which we then averaged across batches and across runs. To maximize statistical power, this final cross-temporal matrix was then symmetrized by averaging the correlation values across reciprocal time point pairs (i.e., averaging the upper and lower triangles of the matrix).

Statistical significance was assessed with a Monte Carlo procedure that preserves the internal structure of each batch. For each batch and time-bin, we created a mismatched odd-session RDM by randomly permuting the 16 stimulus indices within that batch (All permutations were strictly within-batch; stimuli were never mixed across batches). We recomputed the correlation between the permuted odd-session RDM and the even-session RDM. The difference between the real correlation and this permuted correlation was then randomly sign-flipped for each batch. Repeating this 2000 times per run produced a null distribution of geometry consistency values. One-sided p-values were computed as the proportion of null values greater than or equal to the observed value. We applied the Benjamini-Hochberg false discovery rate correction (alpha = 0.05) across time-bins for the time-course and across the upper triangle of the cross-temporal matrix for the time-by-time analysis. In addition to examining each time-bin and each matrix cell, we also wanted to summarize how strongly the early geometry generalizes to the late geometry. This served an important purpose: if early and late geometry appear uncorrelated, this could reflect a true change in representational structure, but it could also reflect a lack of statistical power when studying fine-grained 10 ms bins or performing many comparisons. To rule out the latter explanation, we defined an early window (50 to 90 ms) and a late window (100 to 200 ms) and computed a single summary measure of early-to-late geometry consistency. For each run, we averaged the cross-temporal correlations across all pairs of bins belonging to the early-by-late block. We then applied the same averaging to each surrogate cross-temporal matrix, which yielded a null distribution of early-late values. The p-value was the proportion of null values greater than or equal to the observed mean. This summary analysis provides a sensitive test of whether the early geometry truly fails to generalize to the late geometry and therefore confirms that the two periods reflect distinct representational structures rather than an artefact of temporal resolution or multiple comparisons.

### LSF image construction

To probe whether early vlPFC responses corresponded to a low spatial frequency (LSF) input, we constructed a version of the image set containing only the spatial frequencies below 0.5 cycles per degree (cpd). This corresponds to the spatial frequency evoking the earliest responses in the superior colliculus (Chen et al., 2018) which is receiving direct inputs from the retina and the closest to vlPFC in terms of synaptic relays. To obtain the blurred images, each stimulus was low-pass filtered with an isotropic gaussian. The standard deviation of the gaussian filter was chosen so that the filter amplitude at the cut-off equalled -20 dB.

### DNN features extraction and RSA with early and late geometries

We compared vlPFC representational geometries to a broad panel of 60 pretrained deep neural networks (DNNs), encompassing a wide variety of architectures and training regimes: 20 supervised models, 20 self-supervised models, and 20 language-aligned vision encoders. A complete list of the specific models is provided in Figure S2. For each network, we extracted features from the penultimate layer prior to the final classification or projection head. Specifically, for convolutional architectures, we applied global average pooling to the final spatial feature maps or extracted activations directly from the pre-classification fully connected layer; for Vision Transformers, we extracted the class (CLS) token activation from the final normalization layer; and for language-aligned models, we utilized the visual encoder’s terminal attention pool or layer normalization outputs. RSA analyses were restricted to the 2560 high-variation stimuli (Fig. 1D). The full stimulus set was partitioned into non-overlapping batches of 16 images. Within each batch, euclidean distance-based RDMs were computed for the neuronal data in each time window and the DNN features for ‘Original’ and LSF inputs. For each batch a RSA value was computed with Spearman correlations between the neural and DNN RDMs. For each network × window × image-condition, RSA values were averaged across batches to produce a mean and a SEM. For each time window, we tested whether the mean RSA across DNNs was significantly greater than zero for the Original and LSF conditions using a one-sample sign-flip permutation test of the network-level mean RSA values (10,000 iterations). We tested whether the RSA differed between the two image conditions (Original vs. LSF) using a paired Wilcoxon signed-rank test. The *p*-values from these one-sample and paired tests were combined and corrected for multiple comparisons using the Benjamini-Hochberg false discovery rate (FDR) procedure (ɑ = 0.05). To compare model families, we grouped the networks by training regime (supervised, self-supervised, and language-aligned) and compared their RSA scores for the dominant image condition in each time window using a Kruskal-Wallis H-test. Significant main effects were followed up with pairwise Mann-Whitney U tests, with *p*-values corrected for multiple comparisons using the FDR procedure.

The visualization in figures 1*D* and 1*E* were obtained by applying 2D MDS on RDMs averaged across language-aligned networks processing a random batch of 16 images either in their LSF or original version.

### Pairwise category decoding (from DNN features)

For each of the 28 between-category pairs, we decoded membership of each image from LSF embeddings using logistic regression with a leave-one-sub-category-per-category-out cross validation approach. First, for each DNN’s penultimate layer activation, a PCA (50 components) was fit outside the two categories being decoded and then used to project the pair’s data. Then, for each fold, the classifier was trained on all remaining sub-categories from the two categories and tested on the left-out sub-category pair. For each stimulus, we averaged the classifier’s predictive probability of the true class across the 60 DNNs (see Fig. S2 for individual network performance). Accuracy for the pair was the fraction of images with mean probability ≥ 0.5 for the correct class. We used binomial tests to evaluate whether accuracy values exceeded significantly the chance level. The p-values were corrected across all pairs using the Benjamini-Hochberg FDR procedure.

### Object position decoding from DNN features

The true polar angle of the object centroid was computed from the image metadata (centroid x/y relative to image center). For each network, we predicted angle via linear regression of the penultimate layer feature activation on the 2D target [cosθ,sinθ] with category-leave-out cross-validation: for each category, a PCA (50 components) was fitted on all seven other categories and a linear regressor trained on the 50 dimensional vectors corresponding to each exemplar; held-out category exemplars were then projected and predicted. Per trial, predicted angles were averaged across networks by vector averaging of (cos,sin) (individual network results are provided in Fig. S2). We quantified error as absolute circular difference between predicted and true angles and summarized performance by the MAE across trials. Significance was assessed with a permutation test that shuffled true angles within categories (10,000 iterations), thereby preserving each category’s angle distribution; the p-value was the fraction MAEs from the null distribution that were less or equal than the observed MAE.

### Object position decoding and category decoding from vlPFC data

Analyses were identical to the DNN-based decoding except for the input features. We used vlPFC population vectors built by concatenating all 96 channels across 10-ms bins within the early (50 to 90 ms) or late (100 to 200 ms) windows. For each cross-validation fold and each window, we reduced dimensionality to 50 principal components by fitting PCA on the training data only and then projecting the held-out data through the learned transformation. For position decoding, we used the same category-leave-out scheme: per held-out category, a PCA (50 components) fit on the remaining categories projected the neural vectors, followed by linear regression on [cos(θ), sin(θ)]; errors were summarized by mean absolute circular error (MAE) and tested against a within-category permutation null (10,000 shuffles). For category decoding, we applied the same leave-one-sub-category-per-category-out logistic regression with a global PCA (50 components) fit outside the decoded pair; per-pair p-values from binomial tests were corrected across all pairs using Benjamini–Hochberg FDR.

### Distance to prototype (DP)

To quantify object or sub-category specificity, we measured the distance to prototype (DP) separately in the early (50 to 90 ms) and late (100 to 200 ms) windows. Neural features were concatenated across the 96 channels and the 10 ms bins of the window. For each window we (1) centered the features, (2) performed a PCA to 50 components, and (3) defined, for every sub-category, a prototype as the mean PCA vector using the 10 low-variation images. The DP for that sub-category was the mean squared euclidean distance between the prototype and the responses to its 40 high-variation exemplars. We averaged DP across sub-categories to obtain a window-specific grand mean. To test whether responses were closer to their own prototype than expected from category membership alone, we constructed a category-matched null: for each sub-category we kept its prototype fixed and repeatedly replaced its high-variation set with an equal number of high-variation images from other sub-categories of the same category (10,000 bootstrap resamples per sub-category). The null DPs were averaged across sub-categories, yielding one null value per resample. One-sided p-values were the proportion of null means ≤ the observed mean. Finally, p-values from the two windows were corrected for multiple comparisons using the Benjamini-Hochberg FDR procedure.

### LSF prior “gain” to late window DP

To assess whether coarse, LSF information draws late vlPFC responses toward object prototypes, we compared the mean DP between two sets of exemplars within each sub-category. Specifically, for each sub-category we drew an even split between exemplars based on the category information score returned by DNN classifiers processing LSF images. The category information score was defined as the mean correct-class predictive probability pooled across pairwise low-pass DNN decoders. ΔDP values were then averaged across the 64 subcategories. Significance was assessed with a sign-flip permutation test across objects (10,000 iterations; one-sided, testing for ΔDP < 0).

For the control in Fig. S1 we performed the same procedure but splitting exemplars based on the position information defined as the angular localization error averaged across DNNs. The two p-values testing position information gain and category information gain were corrected for multiple comparisons using the Benjamini-Hochberg FDR procedure.

### RSA feature spaces for faces

Analyses were restricted to face images in the high-variation set. We computed the following RDM between each of the 320 face images: (1) a position RDM object based on the cosine distance between polar angle of the object centroid, (2) a size RDM using euclidean distance between the image scale parameter, and (3) an orientation RDM using the cosine distance between the triplet of rotation value parameters. For each feature and time window, we computed the Spearman correlation between the neuronal euclidean RDM and the position, size and orientation RDMs. We estimated a null distribution (10,000 permutations) by randomly shuffling the feature labels and recomputing features RDM and the correlations with the neural data for which trial order were held fixed. One–sided p-values corresponded to the proportion of permuted correlations that were at least as large as the observed correlation.

### Background/scene representation: image inpainting and model RDM

To assess whether the vlPFC geometry encodes background scene information, we selected, for each object, the 10 low–variation images that differed only in their natural scenes. The RSA was computed using Spearman correlation between the neuronal data euclidean RDM and the average RDM obtained from embeddings of language–aligned DNN processing background–only images obtained by cropping the central object and inpainting the resulting mask.

### Inpainting mask (object removal)

For each object we marked the pixels being identical across the 10 images as the object area and then feathered this area with a Gaussian blur (σ = 12 px) to produce a soft mask. The soft mask defined the region to fill (foreground object) while preserving background structure. We then used StableDiffusion-2 Inpainting (diffusers) to generate candidates of inpainted images. To guide the diffusion process we used the following positive prompt: “empty landscape, photorealistic, consistent lighting, high detail, outdoor, natural scene” and the following negative prompt: “object, animal, person, vehicle, table, text, logo, watermark, focal subject, centered object, unrealistic”. Each inpainted image was manually curated and regenerated if the candidate inpainting image contained artifacts.

### Background model RDM

For each sub-category we extracted penultimate-layer embeddings from language-aligned encoders processing the inpainted images and averaged euclidean RDM across models. We then computed the RSA between the neuronal RDM and the model RDM, per object and time window, using Spearman correlation and averaged RSA values across objects to obtain the panel’s observed correlation. The null distribution (10,000) randomly permuted the neuronal image order within each object before recomputing RSA.

### Correction for multiple comparisons in conscious dimensions analysis

The 8 p-values corresponding to RSA against orientation, size, position, and background models in the early and late windows of Figure 4 were corrected for multiple comparisons using Benjamini-Hochberg FDR.

## Acknowledgments

We thank Ha Hong for providing guidance regarding the use of the image dataset and its metadata. This research was supported by the Templeton World Charity Foundation (Grant numbers 20562, https://doi.org/10.54224/20562 and 33331, https://doi.org/10.54224/33331) to T.I.P. Additional support was provided by the European Union’s Horizon 2020 Framework Programme for Research and Innovation under specific grant agreement no. 945539 (Human Brain Project SGA3). T.v.K. received funding from the European Research Council (ERC) under the European Union’s Horizon 2020 research and innovation program (grant agreement numbers 101078667).

## Author contribution

J.B., T.I.P., and T.v.K. designed research; B.J., T.I.P., and T.v.K. performed surgical procedures; J.B. performed experiments and analyzed data; J.B., M.S., B.J., S.D., T.I.P., and T.v.K interpreted results and wrote the paper; B.J., T.I.P., and T.v.K. supervised research and provided critical revisions; all authors approved the final manuscript.

## Supplementary information

### Supplementary Results

#### Controlling evoked eye movements

In this no-report study the results presented so far cannot be attributed to overt response behavior. Yet, the monkey may have covertly reacted to certain image features by means of pupil size change, microsaccades or eye drifts. We tested whether the eye position, the pupil size and the direction of microsaccades were more consistent between two occurrences of the same stimulus than by chance. Split-half reliability revealed that both eye position and microsaccade direction developed significant stimulus-locked structure. Starting from 160 ms after a stimulus onset, microsaccade direction and eye position tended to be informative about the stimulus (Fig. S3A, left panel).

We next repeated the reliability analysis on a post-hoc microsaccade-free dataset that excluded any trial with a microsaccade onset at or after 100 ms and trials showing residual drifts larger than 0.2° between 100 and 200 ms relative to stimulus onset. As expected by design, no significant reliability remained for eye position (Fig. S3A, right). We then re-ran all core analyses on this ms-free subset. The temporal organization of representational geometry was preserved: early and late windows remained dissociable in time-generalization analyses (Fig. S3B and C). The double dissociation in RSA with deep networks also held: early responses correlated more with LSF representations, whereas late responses correlated more with representations of the Original images (Fig. S3D). Decoding of object position and pairwise category membership remained significant in both windows (Fig. S3E and F), sub-category specificity, quantified as distance-to-prototype, was again absent in the early window and present in the late window (Fig. S3G). Moreover, late-window sub-category convergence scaled with the strength of the low-spatial-frequency category prior, whereas the position information showed no reliable effect (Fig. S3H-I). Finally, the late geometry continued to reflect perceptual dimensions beyond category: it correlated with face orientation, size, and position, and with background-scene structure derived from inpainted images; in contrast, the early geometry reflected face position and background information (Fig. S3J-M). Together, these controls demonstrate that the main results are not a consequence of stimulus-evoked eye movements.

## Supplementary methods

### Oculomotor signal consistency

We measured the potential stimulus-induced idiosyncrasies in ocular signals in bins of 10 ms in the period 0 to 200 ms relative to stimulus onset. For each stimulus, we computed session-wise averages of horizontal/vertical eye position, microsaccade direction and pupil size separately for same odd- and even-indexed sessions used to split the neuronal data in previous analysis. We then computed the time course of between-fold mismatch in eye position as the euclidean distance between odd-session and even-session mean gaze positions (smaller values indicate greater reliability). For the pupil size time course, we used the mean absolute difference between odd- and even-session averages across stimuli. For estimating microsaccade direction consistency, we first computed, per stimulus and time-bin, the circular mean direction separately for the odd and even halves. Then microsaccades directions were mapped to unit complex vectors and the locking index was defined as the magnitude of their mean, |(Vodd + Veven)/2|, averaged across stimuli. When a given stimulus-time-bin combination did not contain any microsaccade event, it was attributed with the circular mean across stimuli in that time-bin to avoid bias from unequal sampling. To test divergence from chance level, we generated a permutation null by randomly re-pairing stimuli between odd and even halves (1,000 permutations), recomputed each metric, and then expressed the observed time courses as z-scores relative to the null distribution at each time-bin. We then applied a two-sided cluster-based permutation test across time (cluster-forming threshold absolute z greater or equal to 1.96; alpha = 0.05) to identify contiguous significant epochs.

### Post-hoc exclusion of stimulus-evoked eye movements (microsaccade-free subset)

Because stimulus-specific changes in ocular signals emerged after 160 ms relative to stimulus onset, we defined a “microsaccade-free” (ms-free) subset of trials to ensure that all main effects reported in the paper do not depend on evoked eye movements. We first identified trials containing a microsaccade whose onset occurred at or after 100 ms post-stimulus; trials meeting this criterion were excluded from the ms-free subset. To further guard against slow drifts or undetected microsaccades, we computed, for each trial, the displacement of mean gaze between 100 ms and 200 ms and excluded trials whose displacement exceeded 0.2° of visual angle. All remaining neuronal analyses described previously in the paper were then repeated on this ms-free subset, using the same preprocessing, session-splitting, and statistics.

## Supplementary Figures

**Figure S1:**
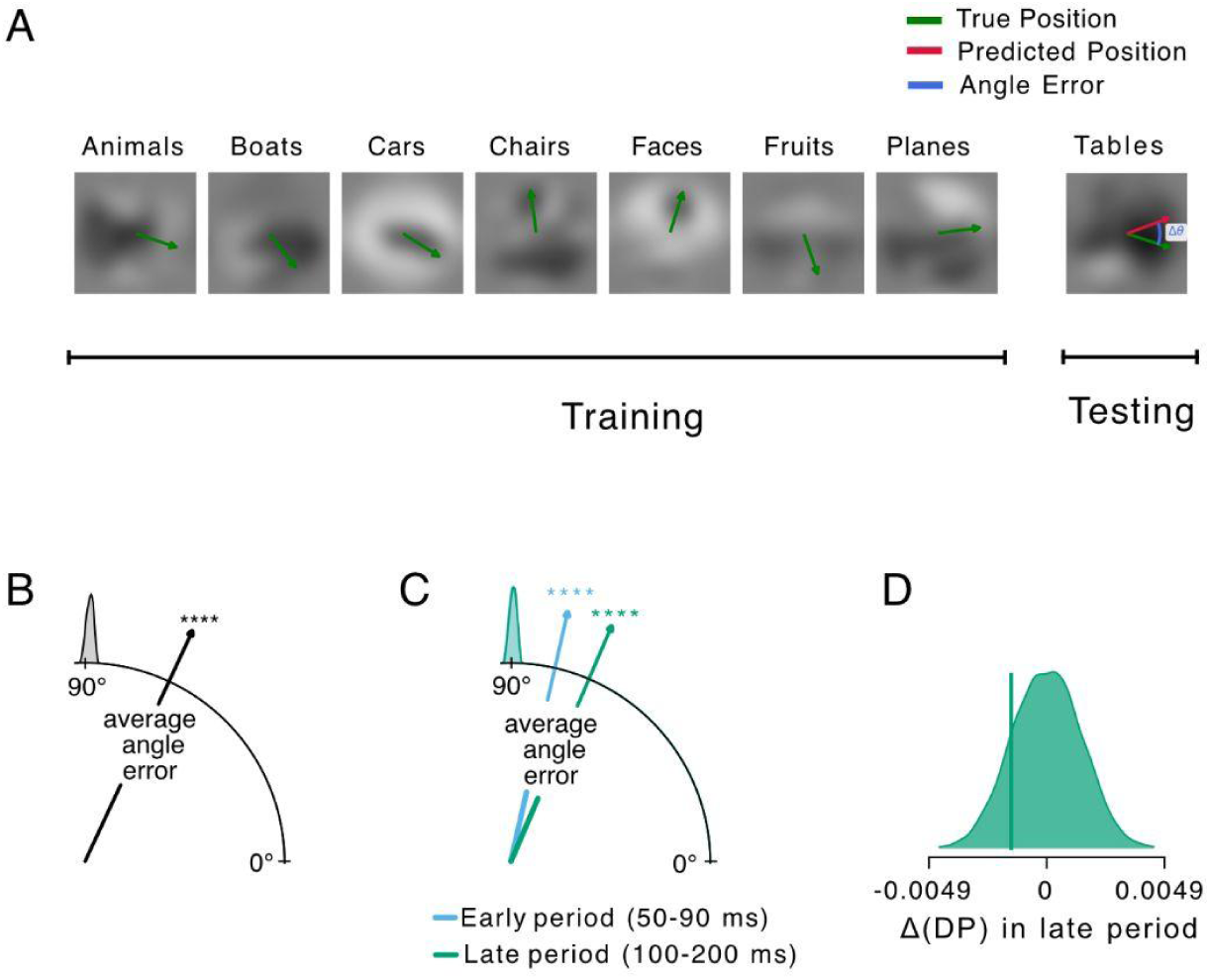
Category-agnostic position information is carried by the LSF component but does not facilitate sub-category information. A. Schematic of the category-leave-out cross-validation strategy for decoding object position. To ensure that position information is abstract and not driven by category-specific features, the decoder is trained to predict the position (green arrow) using images from 7 categories (e.g., Animals, Boats, etc.) and tested on images from the held-out category (e.g., Tables). B. Object position can be decoded from the penultimate layer activation of DNNs processing the blurred images. The arrow is the grand mean of average angle error across the 60 DNNs. The null distribution depicts the surrogate averages obtained by shuffling the position labels across images in the test set. C. Same as B but using the spiking activity from the vlPFC responses. D. Difference in DP between images with low vs. high LSF-based angle error. The DP difference falls within the null distribution, indicating no measurable influence of LSF-derived position estimates on late-phase sub-category discriminability. Note: All face stimuli shown are computer-generated 3D models from the Majaj et al. (2015) dataset and do not depict real human individuals.

**Figure S2:**
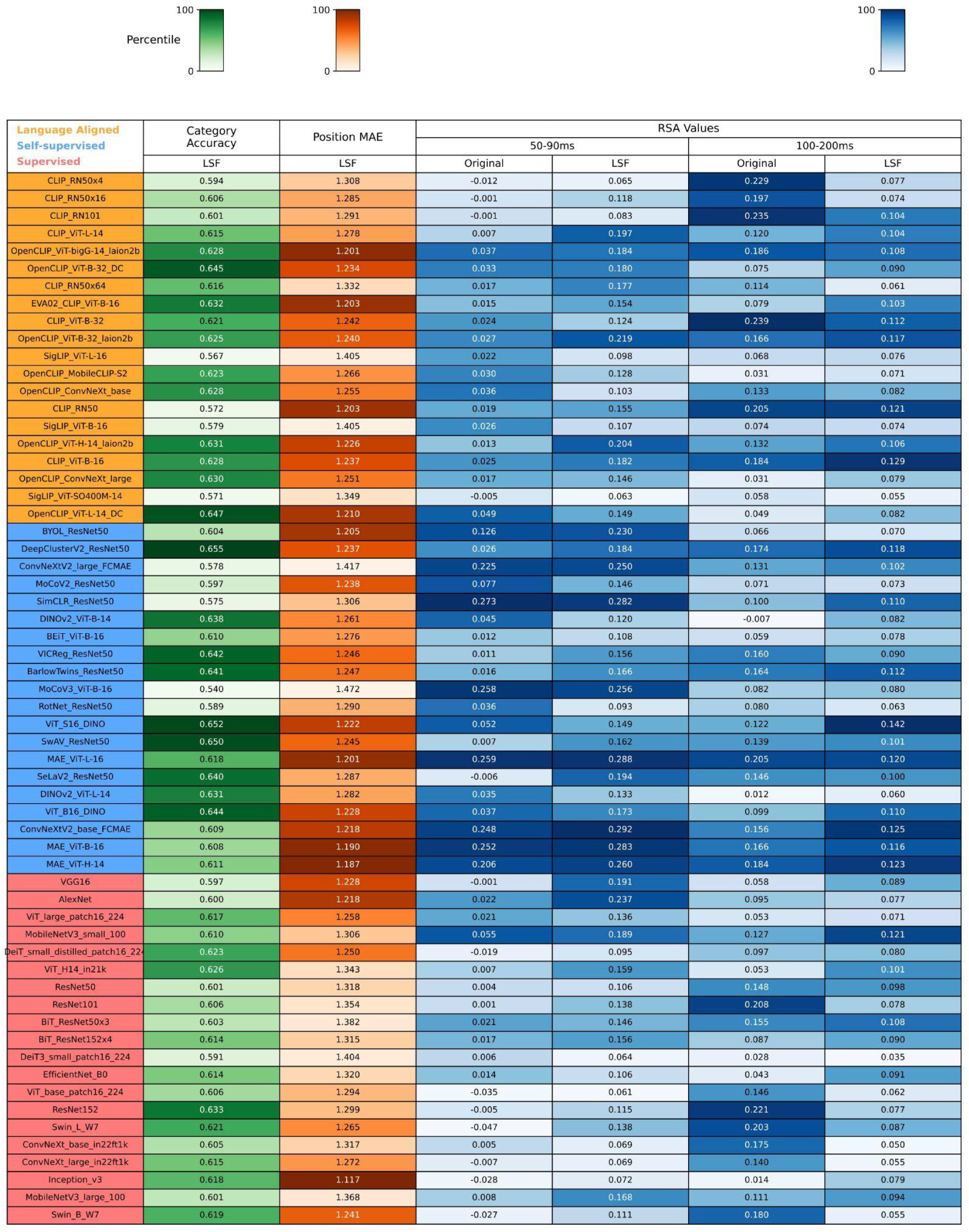
Overview of the performance of each of the 60 DNNs. The first column is the average decoding accuracy for the broad object category from the LSF image. The second column is the Mean Angle Error (MAE) when decoding object position from LSF images (lower is better). The third, fourth, fifth, and sixth columns are the details of the mean RSA values with the vlPFC representations corresponding to Fig. 1F.

**Figure S3:**
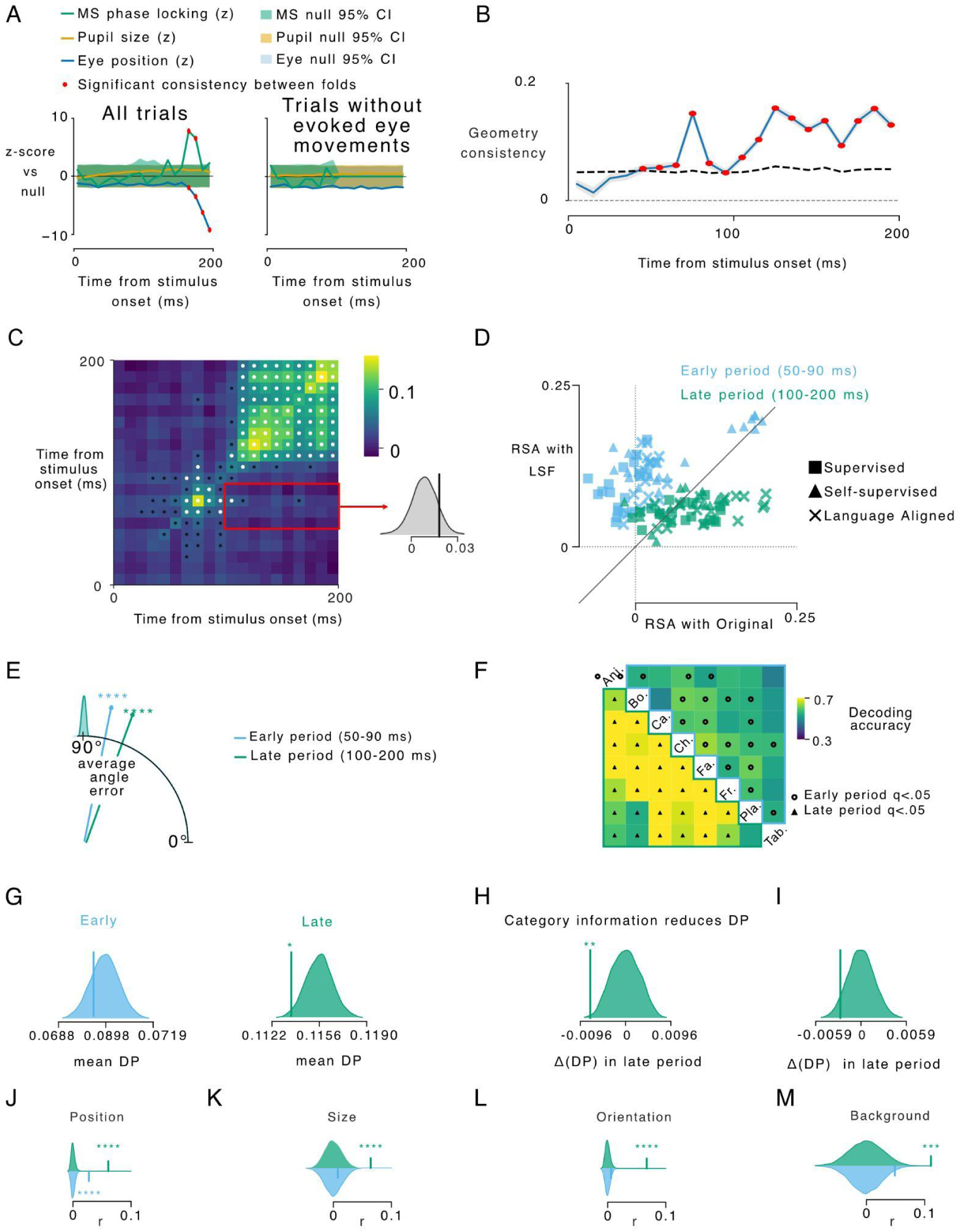
Controlling for stimulus-evoked eye movements. (A) Split-half reliability of oculomotor signals. Left: all trials. Eye position (blue) and microsaccade-direction locking (green) become stimulus-informative only after 160 ms; pupil size (orange) shows no reliability. Shaded bands: 95% permutation-null CI; red dots: cluster-corrected significant bins. Right: microsaccade-free subset excluding trials with microsaccade onset ≥100 ms and trials with residual gaze drift >0.2° between 100 and 200 ms; post-100-ms reliability is abolished and the microsaccade trace is undefined after 100 ms by design. (B) Time-course of geometry consistency. (C) RSA with DNNs. (D) Cross-temporal generalization of representational geometry. (E) Object-position decoding from neural activity. (F) Pairwise category decoding from neural activity. (G) Distance-to-prototype (DP) by window. (H). Influence of low-spatial-frequency category priors on late-window DP. (I) Position information control. (J to M). Representational similarity between vlPFC geometry and perceptual features beyond category; stars indicate FDR-corrected significance.

